# Atypical developmental features of cortical thickness trajectories in Autism Spectrum Disorder

**DOI:** 10.1101/580837

**Authors:** Adonay S Nunes, Vasily A Vakorin, Nataliia Kozhemiako, Nicholas Peatfield, Urs Ribary, Sam M Doesburg

**Author notes:** Correspondence Biomedical Physiology Department, Simon Fraser University, 8888 University Dr, Burnaby, BC V5A 1S6, Canada.

## Abstract

Neuroimaging studies have reported numerous region-specific atypicalities in the brains of individuals with Autism Spectrum Disorder (ASD), including alterations in cortical thickness (CT). However, there are many inconsistent findings, and this is probably due to atypical CT developmental trajectories in ASD. To this end, we investigated group differences in terms of shapes of developmental trajectories of CT between ASD and typically developing (TD) populations.

Using the Autism Brain Imaging Data Exchange (ABIDE) repository (releases I and II combined), we investigated atypical shapes of developmental trajectories in ASD using a linear, quadratic and cubic models at various scales of spatial coarseness, and their association with symptomatology using the Autism Diagnostic Observation Schedule (ADOS) scores. These parameters were also used to predict ASD and TD CT development.

While no overall group differences in CT was observed across the entire age range, ASD and TD populations were different in terms of age-related changes. Developmental trajectories of CT in ASD were mostly characterized by decreased cortical thinning during early adolescence and increased thinning at later stages, involving mostly frontal and parietal areas. Such changes were associated with ADOS scores. The curvature of the trajectories estimated from the quadratic model was the most accurate and sensitive measure for detecting ASD. Our findings suggest that under the context of longitudinal changes in brain morphology, robust detection of ASD would require three time points to estimate the curvature of age-related changes.

## Introduction

Autism Spectrum Disorder (ASD) is a highly heterogeneous neurodevelopmental disorder that has an early-life onset. It is characterized by deficits in social communication and restricted or repetitive behaviors and/or interests ^1^. Several neuroimaging studies have reported region-specific changes in brain morphology in ASD ^2–4^, and supported by post-mortem studies ^5^. Findings of atypical brain morphology in ASD, however, are highly heterogeneous ^6–8^, and there are not yet clear neuroanatomical markers for accurately identifying individuals with ASD.

ASD is a very heterogeneous disorder which together with the small sample sizes and varying age ranges of some studies might partially explain the inconsistencies in the neuroimaging literature ^9,10^. One hypothesis is that such inconsistencies are due to the age-specific developmental atypicalities of ASD brain. Many studies have recently supported this, reporting atypical developmental trajectories in the ASD brain in terms of neuroanatomy ^11–13^, hemodynamic functional connectivity ^14–16^ and neurophysiological rhythms and synchrony ^17–19^.

Significant alterations of developmental trajectories of CT have been reported in ASD. For example, it has been shown that in ASD adults CT decreases more dramatically with age compared to TD ^20^. Increased CT in children with ASD has also been reported in children aged 8-12 years ^3^. Zielinski et al. (2104) reported three distinct phases of atypical cortical development in ASD: an accelerated expansion during early childhood, accelerated thinning in later childhood and adolescence and decelerated thinning in early adulthood ^21^. Wallace and colleagues, conversely, reported accelerated age-related cortical thinning in temporal and parietal areas during early adulthood in the ASD population ^22^. In addition, based on a sample of 6-15 year old children with ASD, Jiao and colleagues found both increased and decreased thickening of cortical areas ^23^. Recently, Kundrakpam et al. (2017) using the ABIDE I, reported higher CT in ASD until adolescence where accelerated thinning was found until adulthood.

In the present study, we used the largest cross-sectional database available to capture developmental shapes of CT trajectories that are atypical in ASD between childhood and late adolescence. Age-related changes in cortical thickness have been previously explored based on linear, quadratic, and cubic models ^24–27^. Instead of deriving statistics from fitting a model that describes age-related CT differences between ASD and TD, we used a linear, quadratic and cubic models to extract their highest order derivative. The coefficients of these derivatives are constant across the entire age range and have a geometrical interpretation, herein, the developmental features are referred to as “trajectory shapes”, and the highest order coefficient in the linear model (the linear trend) as the ‘slope’, in the quadratic (acceleration) as the ‘curvature’ and in the cubic (rate at which acceleration changes) as the ‘aberrancy’.

Previously, brain morphometric measures such as CT and cortical volume have been used to predict ASD^23,28–30^, and remains a promising feature. Recently, an ABIDE study used linear developmental changes in white/gray matter contrast to predict ASD development^31^. In the present study, we trained a classifier by either using the slope, curvature or aberrancy to predict ASD development. Although a cross-sectional sample is used, this classification could be applied longitudinally to classify individuals with ASD, and our results could provide evidence of the necessary longitudinal time points to estimate a specific trajectory shape.

We explored atypical trajectory shapes at three levels of coarseness to detect if abnormal CT maturation is generalizable across the cortex or if it is spatially very constrained to specific areas. We used three atlases. The first atlas is a trivial one, simply representing two hemispheres. The second atlas is based on a typical FreeSurfer anatomical parcellation (FSAP): 70 brain areas. Finally, the third atlas is based on the Multi-Modal Parcellation^32^ (MMP): 360 brain areas. We will refer to these three parcellations as hemispheric, anatomical, and multi-modal, respectively, representing a spectrum of spatial coarseness from the most coarse to fine-grained. In addition, the trajectory shapes (the slope, curvature and aberrancy) obtained at each area of the FSAP and MMP were correlated with ADOS scores to assess if symptom severity contributes to more exacerbated atypical trajectory shapes in ASD.

## Results

### Overall CT is not atypical in ASD when averaged across a large age range

First, we assessed group differences in CT between experimental groups at the hemispheric, anatomical and multi-modal levels. The CT across sites were corrected for different scan parameters, as illustrated in Fig. 1 step 6. Statistical significance was tested using the multivariate mean-centered PLS analysis (see methods). It revealed no overall group differences in CT between ASD and TD groups for all the atlases. This can be clearly seen in the CT trajectories (Fig. 2 & 4) where differences are marked by changes across age rather than differences in mean CT between the two groups.

**Fig 1.**
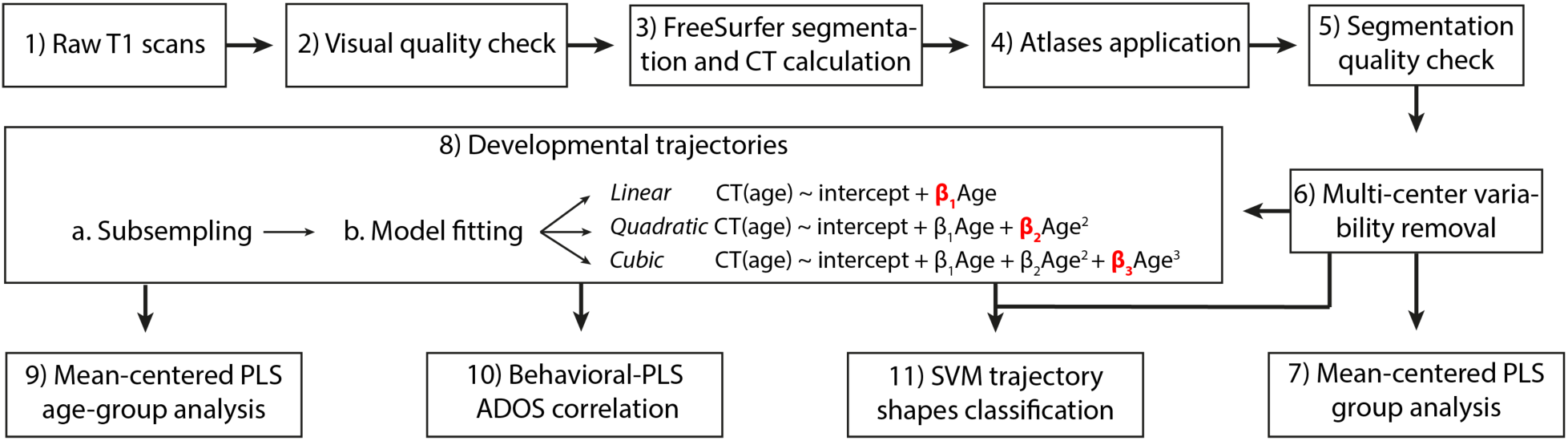
Workflow diagram of the steps of this study. The coefficients in red in step 8 are the trajectory shapes used in the subsampling, correlation and classification analyses.

**Figure 2.**
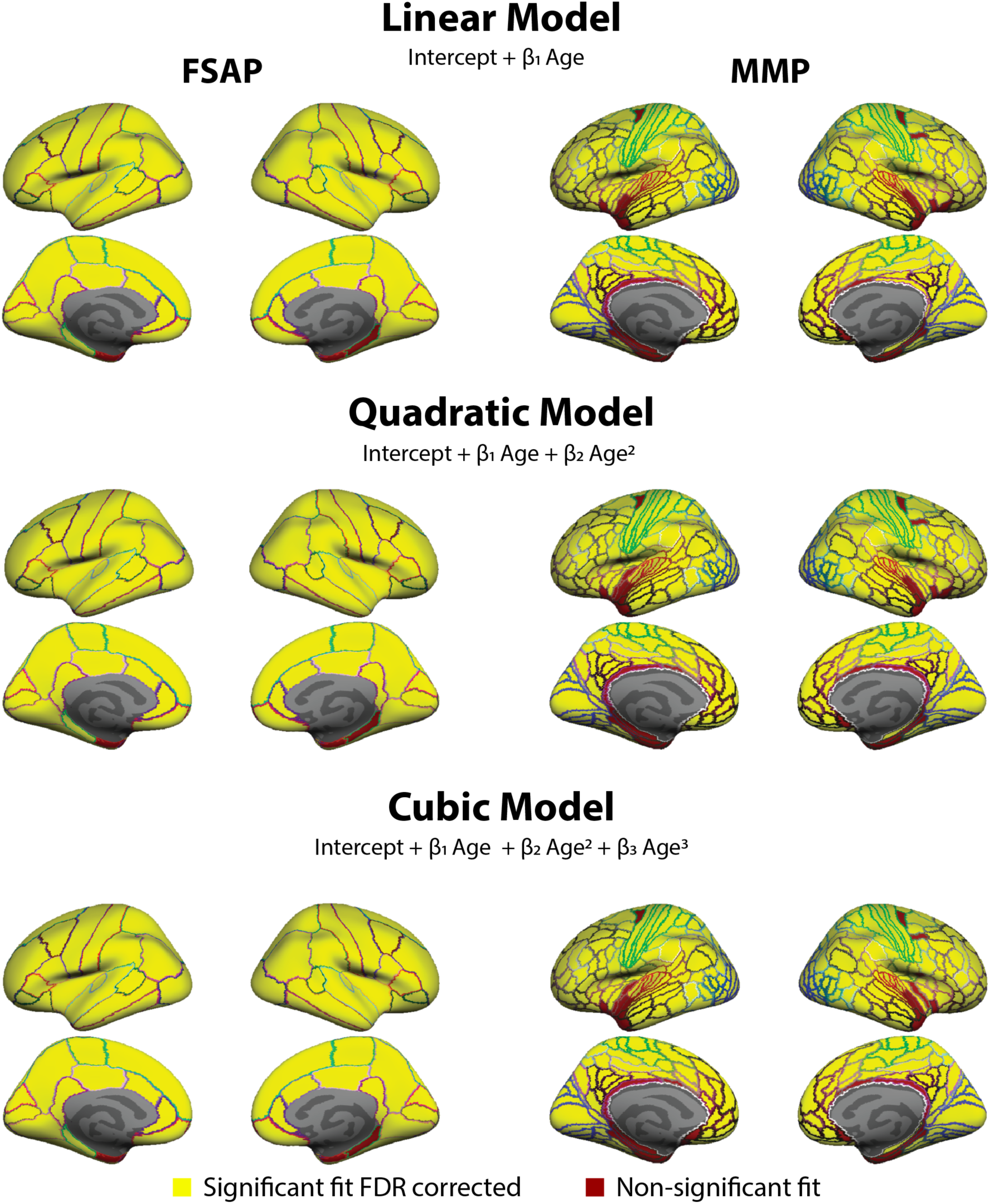
Areas from the FSAP and MMP used in the analyses. The goodness of fit at each area of the FSAP and MMP was measured with a deviance test assessing if the model significantly fits better than a constant model. After FDR correction, areas with a significant fit were used in subsequent analyses (colored in yellow), and non-significant were masked out (colored in red). Colored areal border outlines represent the original atlas annotation color of the FSAP (left) and MMP (right).

### Atypical developmental trajectories of CT in ASD

Linear, quadratic and cubic models were fitted, separately for each group, to estimate CT trajectory shapes in each area of the three parcellations. First, we fitted the models using all the subjects from a group to test the goodness of fit in a given area, then, from subgroups derived using a subsampling technique to extract trajectory shapes. On the group level fitting, the goodness of fit of a model in each area was tested with a deviance test. Illustrated in Fig 2, a few areas around the ventral medial temporal cortex, models did not significantly fit, especially at the multi-modal level using the MMP. These areas were masked out and excluded from further analyses.

Using the subsampling approach, in each group we generated 80 subgroups composed of 70 subjects and the three models were fit. From each area of the parcellations, we extracted a trajectory shape from each model (slope, curvature and aberrancy). Then, these trajectory shapes were statistically assessed using mean-centered PLS. We found statistically significant differences in developmental trajectory shapes in CT for the hemispheric, anatomical and multimodal parcellations rendering a p-value of p<0.01, p<0.001 and p<0.001 respectively.

At the hemispheric level, the statistical analysis indicated strong effects in the quadratic model with a curvature value more negative (less concaved) in the ASD, in the linear model with the slope being more positive (slower decline), while the aberrancy of the cubic model did not contribute much to the hemispheric CT group differences. These effects can be seen in Fig. 3 with the z-scores for each model plotted on the brain surfaces. The curvature coefficient has a z-score of -11 for both hemispheres, the linear 3 and cubic 0. The effects captured by the curvature and the slope are very similar but with an opposite sign. In this case, the effect reflected in the slope indicates decreased thinning in the ASD, while the curvature indicates decreased thinning, compared to TD, until around 15 years of age and then accelerated thinning. The aberrancy did not capture stable effects.

**Figure 3.**
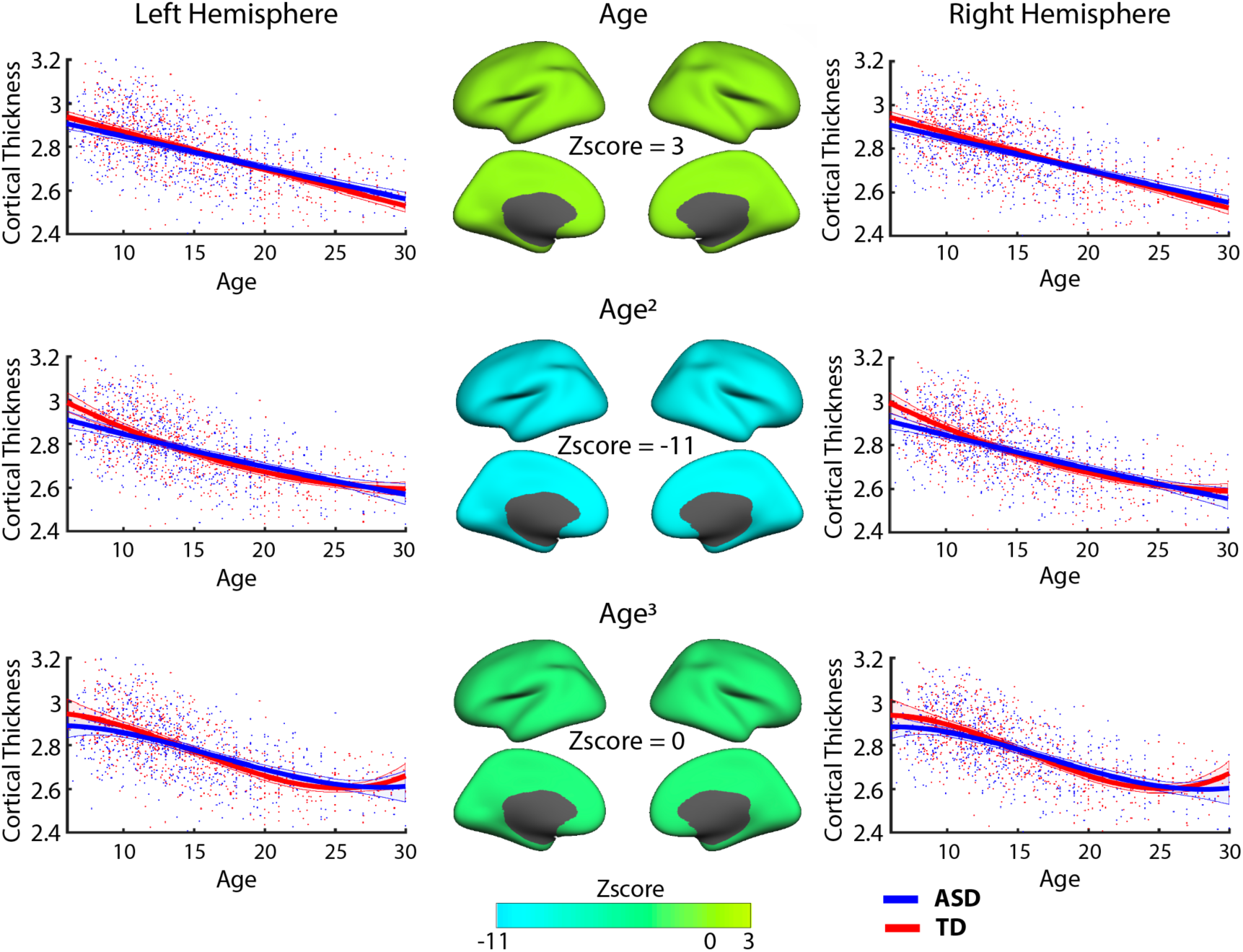
Developmental trajectories of hemispheric CT. The slope, curvature and aberrancy parameters from the three models were used to find altered trajectory shapes in ASD. In the middle, z-scores from the PLS analysis are plotted for each hemisphere representing how reliable the parameters were in expressing the overall group differences. The trajectory shape represented by the curvature coefficient was the strongest followed by the slope.

For the anatomical and multi-modal parcellations, mean-centered PLS analyses rendered similar results as the hemispheric, and the multi-modal was spatially more defined. As in the hemispheric level, the results between the linear and quadratic model were spatially similar but with opposite sign z-scores. In Fig. 4, z-scores for each model and atlas are visualized. The z-scores are with respect to the contrast ASD>TD. The areas with the highest z-scores representing the most stable group differences were located in frontal, fontal-medial and temporoparietal junction (TPJ) areas for the linear and quadratic models. In addition, for the quadratic model, brain regions with high z-scores were localized in the posterior cingulate cortex (PCC). In contrast, z-scores for the cubic model were located in occipital and posterior-parietal areas. Fig. 5 illustrates the developmental trajectories from areas with high z-scores for a given model.

**Figure 4.**
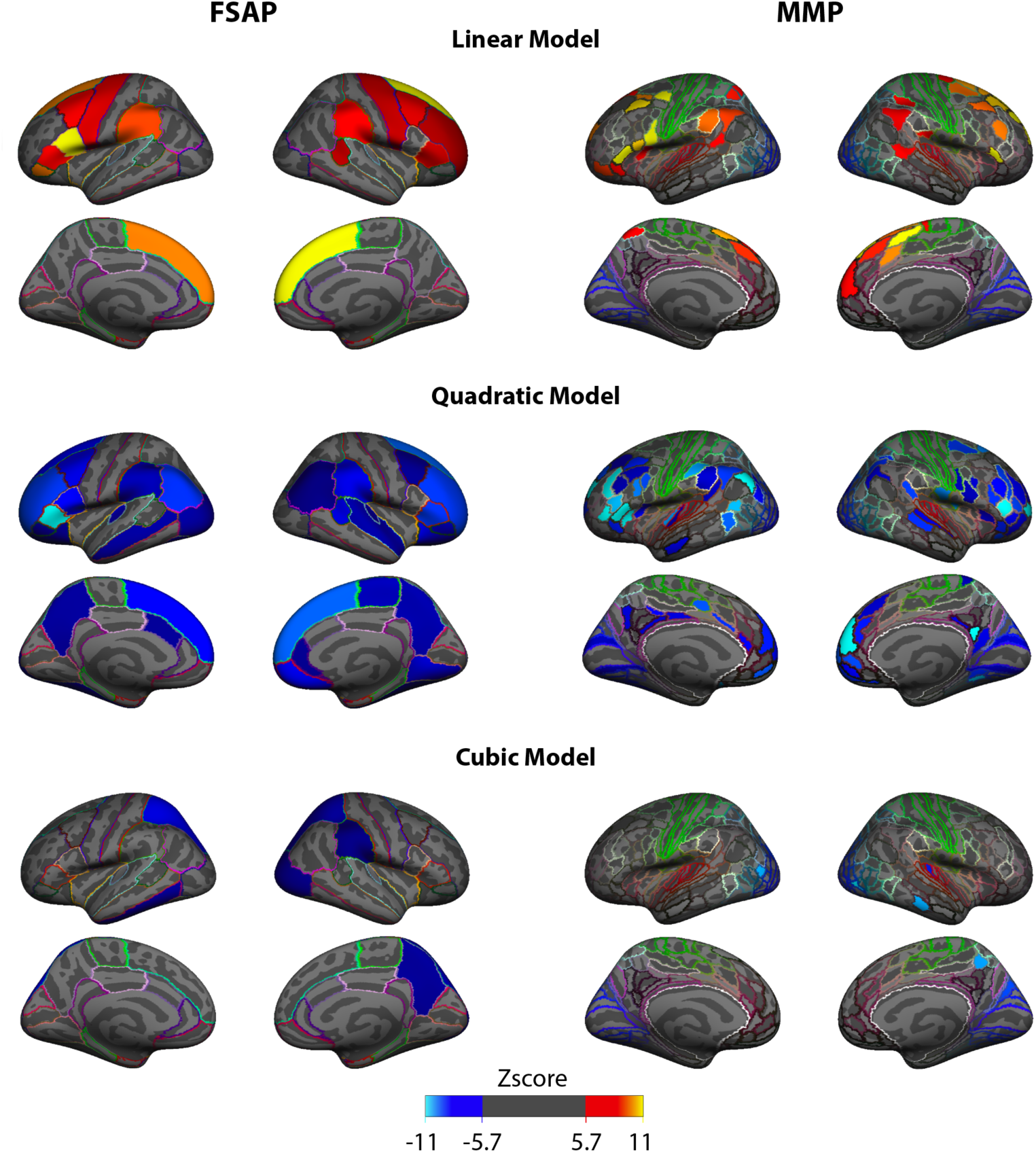
Distributions of the z-scores across brain areas, shown separately for three models and two atlases (left for anatomical, right for multi-modal). For the group contrast (ASD>TD) the effect found in the linear coefficient is positive, whereas in the quadratic and cubic is negative. The spatial distribution of the highest z-scores in the linear model is very similar to the quadratic, and both represent a decreased cortical thinning during childhood but the quadratic also captures accelerated thinning after mid-adolescence. Colored border outlines in each cortical area represent the original atlas annotation color of the area.

**Figure 5.**
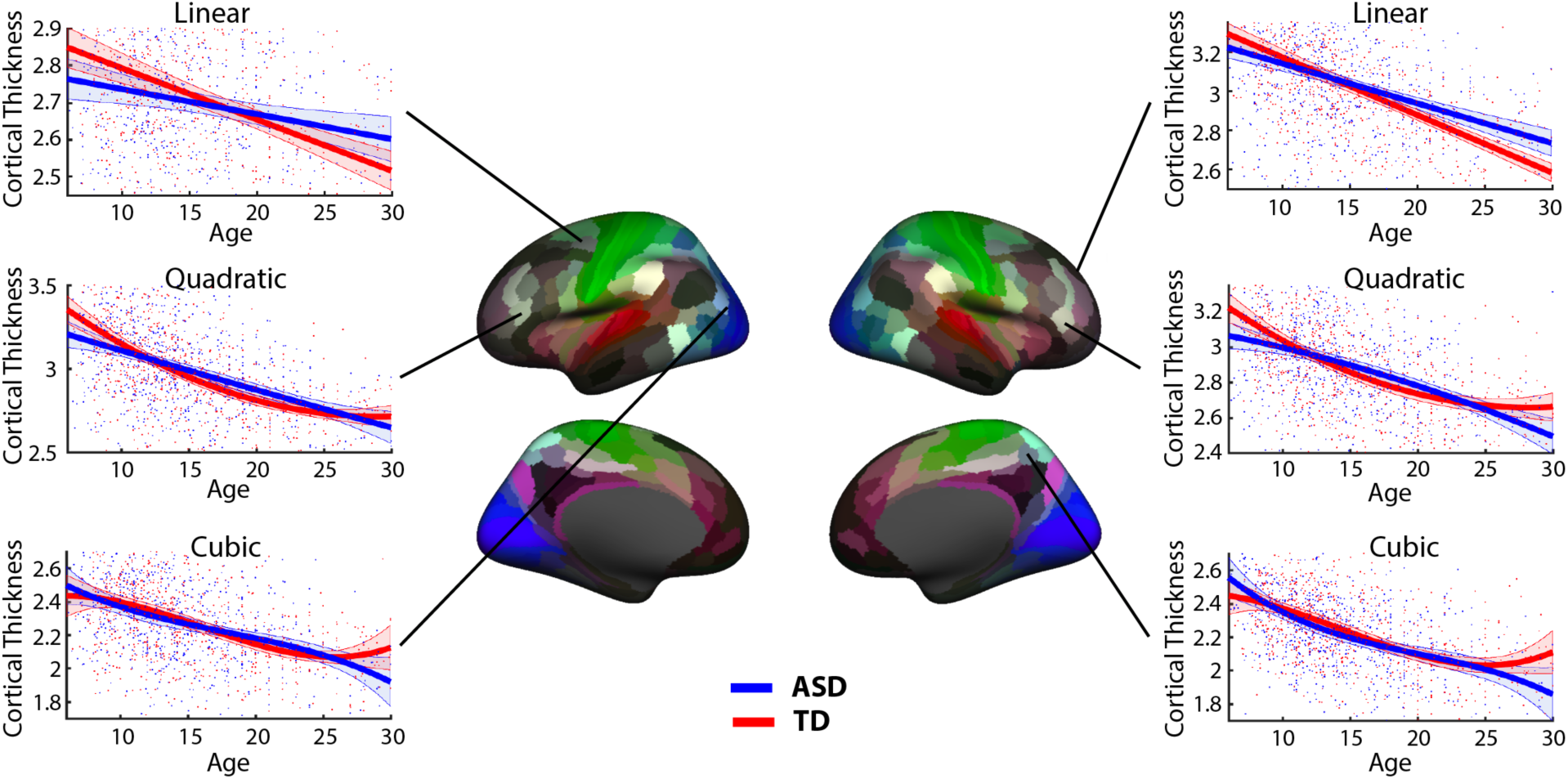
Developmental trajectories in areas that reliably expressed group differences. Trajectories are plotted selecting areas with high z-scores values for the right and left hemisphere for the three models used.

### Center-specific analysis of developmental trajectories of CT in ASD

To further support our findings based on data in which batch effects (multi-center variability) were removed, the same analysis was performed using subjects from only one center. As in the results from the full dataset, the z-score distribution for the linear coefficient was also skewed on the right and for the curvature coefficient on the left. The same effect of decreased thinning at an early age (captured by the linear and quadratic) and acceleration during mid-adolescence (captured by the quadratic) were found with the NYU dataset. As can be seen in Supp. Figure 1, areas with the highest z-scores for the linear and quadratic coefficients are also located in frontal and temporal lobes, TPJ and PCC. The effects captured by the cubic coefficient did not correspond between the full and center-specific analyses, suggesting once more that the aberrancy of the cubic model did not capture stable effects.

### Associations between ADOS and developmental trajectories of CT

To explore the relationship between ASD symptomatology and trajectory shapes in CT maturation, behavioral PLS correlated the trajectory shape of a model fitted in a subgroup with the subgroup mean ADOS-Generic scores (communication, social and stereotyped behavior subscales). This measured if the subgroup’s trajectory shape is associated with ADOS scores. Given that the trajectory shapes might correlate differently, for each of the three trajectory shapes at the anatomical and multi-modal spatial levels we performed different behavioral-PLS (6 in total), and the p-values obtained for each model were Bonferroni corrected.

Only the overall correlations between the curvature of the quadratic model from the FSAP and MMP and ADOS scores were significant after Bonferroni correction with a corrected p-val <0.01 for both levels. The distribution of z-scores for both parcellations was skewed to the left, which indicate negative correlations between the curvature values and ADOS. The more negative the curvature, which in this case represents accelerated CT thinning after mid-adolescence, the higher the symptom severity. More specifically, the correlation values between the curvature coefficients from the FSAP and ADOS sub-scores were: r= -.21 for communication subscale, r= -.22 for social subscale and r= -.11 for stereotyped behavior subscale. For the MMPA the correlations were: r= -.27 for communication subscale, r= -.27 for social subscale and r= -.16 for stereotyped behavior subscale. Fig. 6 illustrates the LV design reflecting the contribution of each ADOS sub-score, their correlation values with curvature coefficients, and the spatial topography of the most negative z-scores. The spatial topography of the most negative z-scores was very similar to the group differences analysis, being mostly located in frontal areas, TPJ and PCC.

**Figure 6.**
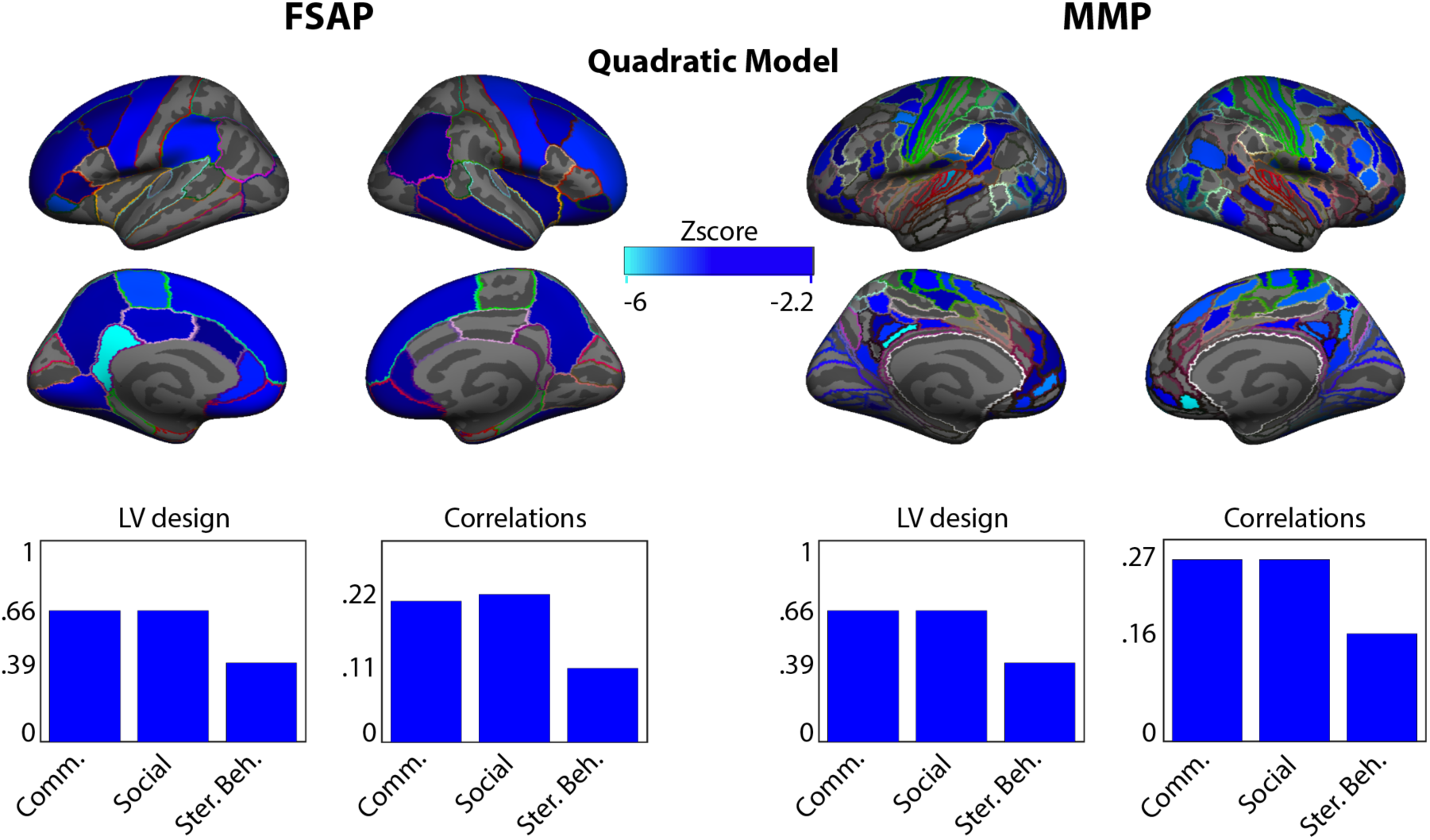
Correlations between the ASD curvature values and mean ADOS scores averaged across the subjects used to characterize the curvature. The spatial map in blue indicates negative z-scores representing the association between ADOS scores and a negative curvature developmental trajectory. On the bottom, raw histograms are plotted for each atlas representing the contribution of the ADOS scores to the latent variable (how much each contributed to the overall correlation) and the correlation of each ADOS score with all the curvature coefficients of the areas.

### ASD classification of CT developmental trajectories

A support vector machine was used to classify trajectory shapes in CT from ASD and TD subgroups. These trajectory shapes reflect developmental features that could be obtained longitudinally from a single subject, as their derivative is constant across age. For the anatomical and multi-modal parcellations, using the FSAP and MMP respectively, an SVM was trained to classify trajectory shapes from the linear, quadratic and cubic highest order coefficients. Figure 7 illustrates the distribution of the accuracy, specificity and sensitivity obtained with 500 cross-validation runs.

**Figure 7.**
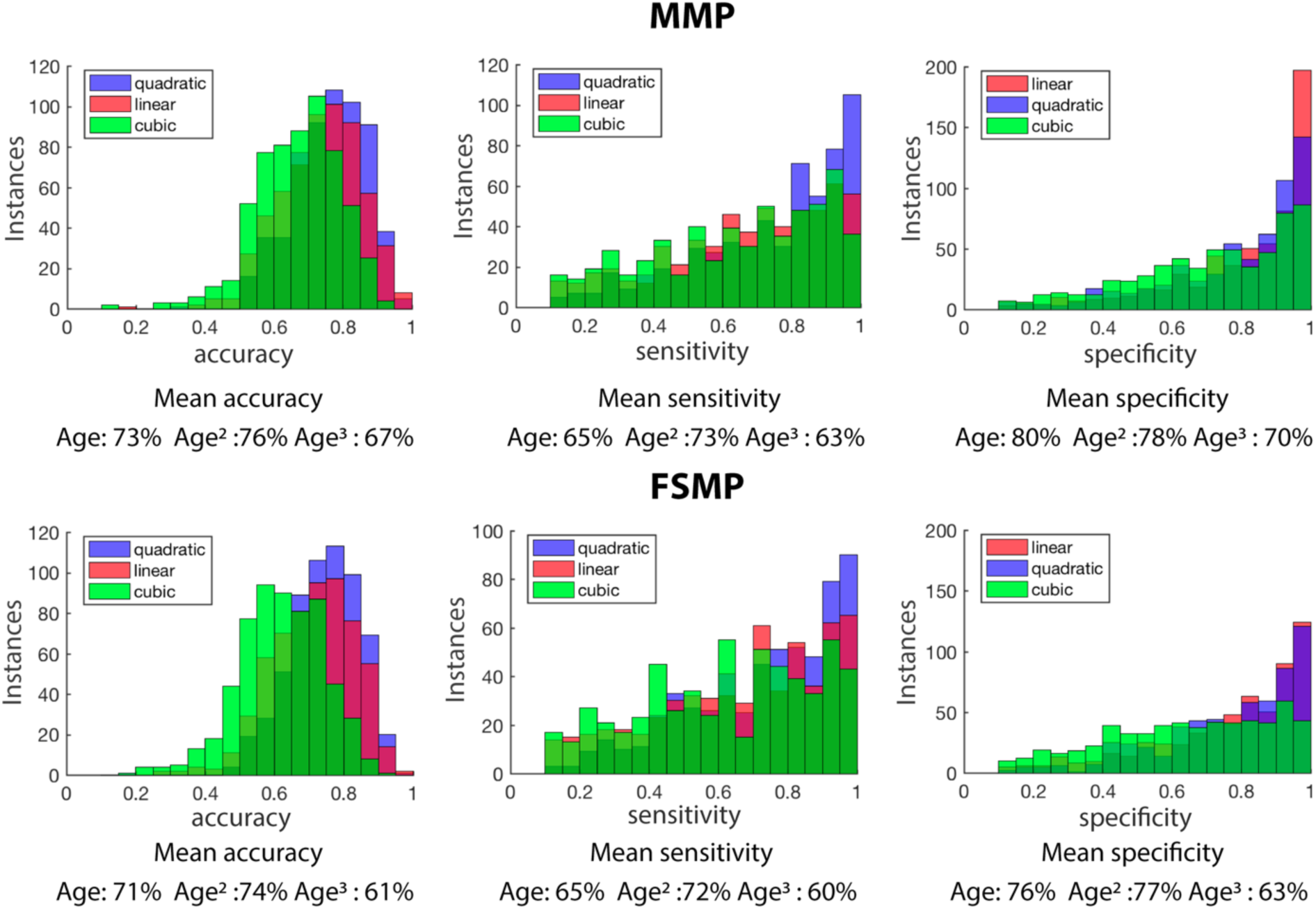
Distributions of classification accuracy, sensitivity and specificity using an SVM to classify ASD and TD trajectory features of CT development using areas from the MMP and FSMP. Each instance was a cross-validation run where the dataset was randomly split into two equal halves for training and testing.

The best total accuracy in classifying ASD and TDC was archived using the quadratic trajectory shape with an accuracy of 76% with the MMP and 74% with the FSAM, followed by the linear (73% and 71%) and cubic (67% and 61%). When separating accuracy in sensitivity (ASD detection) and specificity (CTR detection), using the MMP the curvature identified more ASD (mean 73%), while the slope more CTR (80%).

## Methods

### Participants

We analyzed structural T1-weighted MRI scans from the Autism Brain Imaging Data Exchange (ABIDE) repository (releases I and II). Quality control was assessed by visual inspection by three independent reviewers, and subjects with strong movement artifacts were discarded. To reduce the bias in the age distribution, we excluded participants younger than 6 years of age and older than 30 years of age as beyond this age range the number of participants drops significantly. To consider the full spectrum of ASD cohort, we did not exclude subjects based on their IQ.

Out of 2226 subjects available in the ABIDE I & II, 571 were rejected based on motion artifacts detected during the visual inspection or segmentation errors revealed at the preprocessing stage, as explained below. To properly account for center variance, centers with less than 10 subjects in each group were removed, which resulted in 24 included in the analysis. The centers with the same parameters from ABIDE I and II were combined. In total, our final sample included 674 subjects with ASD (females = 94, mean 14.5 ±5.18 years) and 686 TD subjects (females=134, mean 14.9 ±5.49 years). Age between ASD/TD, the proportion of male/female in each center between groups or the proportion of ASD/TD subjects per center did not differ significantly. Information on the participants can be found in Table 1. 404 ASD participants had information on ADOS-Generic scores (communication, social and stereotyped behavior) and were included in the analysis of associations between ADOS and age-related trajectory shapes in CT.

**Table 1.** Description of the number of participants per center, number of females per center, age mean and standard deviation. Full names of the centers available at: http://fcon_1000.projects.nitrc.org/indi/abide/

| Centers | N |  | Females |  | Mean age |  | STD age |  |
| --- | --- | --- | --- | --- | --- | --- | --- | --- |
|  | ASD | TD | ASD | TD | ASD | TD | ASD | TD |
| **CALTECH** | 14 | 12 | 3 | 4 | 22.1 | 22.6 | 3.1 | 3.2 |
| **CMU** | 10 | 10 | 1 | 3 | 23.5 | 24.5 | 3.6 | 3.5 |
| **KKI** | 58 | 61 | 16 | 16 | 10.3 | 10.3 | 1.5 | 1.2 |
| **LEUVEN** | 24 | 25 | 2 | 2 | 17.6 | 18.1 | 4.2 | 4.8 |
| **MAX MUN** | 13 | 13 | 1 | 0 | 18.3 | 22.8 | 8.1 | 6.7 |
| **NYU** | 109 | 116 | 12 | 23 | 12.5 | 13.8 | 5.8 | 5.4 |
| **OHSU** | 30 | 30 | 4 | 6 | 11.2 | 10.6 | 2.1 | 1.7 |
| **OLIN** | 16 | 14 | 3 | 2 | 16.8 | 16.9 | 3.4 | 3.6 |
| **PITT** | 24 | 23 | 4 | 4 | 16.5 | 17.5 | 4.7 | 4.6 |
| **SDSU** | 37 | 41 | 7 | 7 | 13.8 | 13.7 | 3.1 | 2.5 |
| **STANDFORD** | 15 | 17 | 4 | 4 | 10.1 | 9.9 | 1.6 | 1.6 |
| **TRINITY** | 40 | 45 | 0 | 0 | 15.9 | 16.5 | 3.4 | 3.5 |
| **USM** | 37 | 33 | 2 | 3 | 12.9 | 12.1 | 2.4 | 2.5 |
| **YALE** | 24 | 26 | 7 | 8 | 13.5 | 14.6 | 2.6 | 4.0 |
| **UCLA** | 60 | 53 | 7 | 11 | 19.3 | 20.0 | 5.2 | 6.1 |
| **UM** | 42 | 48 | 5 | 8 | 13.0 | 12.9 | 3.0 | 2.7 |
| **BNI** | 11 | 10 | 0 | 0 | 21.1 | 20.8 | 2.2 | 2.6 |
| **EMC** | 14 | 14 | 2 | 2 | 8.5 | 8.1 | 1.3 | 1.1 |
| **ETH** | 10 | 10 | 0 | 0 | 21.1 | 24.0 | 3.5 | 4.0 |
| **GU** | 31 | 31 | 4 | 10 | 11.1 | 11.1 | 1.4 | 1.6 |
| **IP** | 18 | 17 | 5 | 10 | 15.3 | 18.8 | 4.9 | 6.4 |
| **IU** | 15 | 17 | 2 | 5 | 21.5 | 21.9 | 3.5 | 2.0 |
| **ONRC** | 10 | 9 | 1 | 2 | 22.2 | 22.7 | 3.6 | 3.4 |
| **UCD** | 12 | 11 | 2 | 4 | 15.3 | 15.1 | 2.0 | 1.4 |
| **Total** | 674 | 686 | 94 | 134 | 14.5 | 14.9 | 5.2 | 5.5 |

Our analysis was conducted on data from all centers combined, after correcting for batch effects (variability across centers), and, to validate the results excluding the possibility of being driven by batch effects, on a single center using the New York University (NYU) repository which has the largest sample size in the ABIDE dataset.

### Preprocessing, CT calculation and regions of interest

MRI data were segmented using FreeSurfer ^33^. Cortical thickness (CT) was computed as the distance between the white and the pial surfaces for each vertex. CT values were averaged at three levels: hemispheric, anatomical and multimodal. First, CT values were averaged within hemispheres (hemisphere parcellation, HP, with two CT values). Second, at the anatomical level, CT values were averaged within the FreeSurfer Anatomical Parcellation (FSAP, with 68 areas) ^34^. Third, at the multi-modal level, using the MultiModal Parcellation (MMP, with 360 areas) atlas whose areas are defined by sharp changes in cortical architecture, function, connectivity, and/or topography ^35^.

To assess the presence of segmentation errors in reconstructing the cortical mantle from which CT was estimated, we conducted a second quality check. For every center separately, in each parcellation area, we performed a z-score transformation across subjects, resulting in the z-score distribution with zero mean and unit variance for each center. Subjects with a z-score with a magnitude bigger than 3 in any area of the parcellations were considered outliers and excluded from further analysis.

### Inter-center variability removal

Previous studies have reported variability in MRI-based measures across centers in the ABIDE repository (Haar et al., 2014). To correct for center differences, at each area of the parcellations a linear, quadratic and cubic models were fitted to estimate the variance explained by centers while preserving the variance explained by group, age and group x age interactions. Specifically, the following equations were applied:

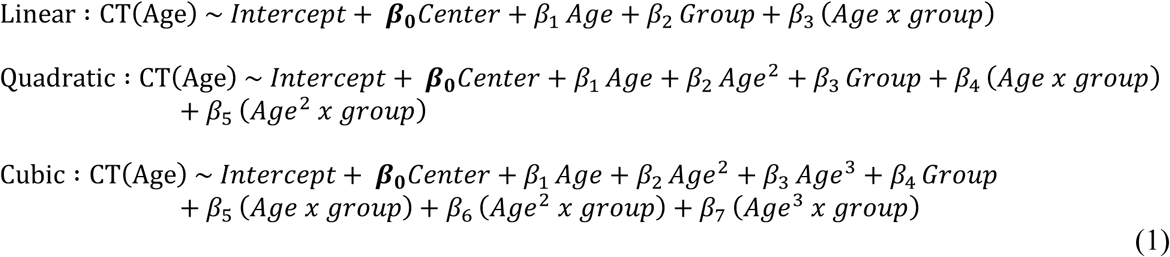

Where *Center* is a binary variable with the size (N subjects × N centers) and the ***β***_**0**_ coefficients in each of the three models explain the center variance for a given area. The variance explained by ***β***_**0**_*Center* was regressed out from each area in the HP, FSAP and MMP. After this center correction, CT values across centers were combined for further analyses.

### Modeling developmental trajectories of cortical thickness

Following the model formulas in Eq.1, from the linear model we obtained the *β*_1_ coefficient (linear trend), or ‘slope’, from the quadratic model we obtained the *β*_2_ coefficient (acceleration) or ‘curvature’, and from the cubic the *β*_3_ coefficient (rate at which acceleration changes) or ‘aberrancy’.

To capture trajectory shapes in each experimental group, we fit the three models separately for ASD and TD for brain areas for each of the three parcellations using the CT data corrected for center variance. Given that models were fitted separately per group, the parameters group and age × group interaction from eq.(1) were not necessary, as well as the center parameter ***β***_**0**_ as the center variance was regressed out in the previous step. Formulas for the current models are depicted in workflow figure (Fig.1, step 8).

First, these models were fit on the entire group sample to test the goodness of fit, and after, on each subsample (or subgroup) to capture the within group variability in trajectory shapes, as explained in the subsample analysis section below.

On the group level fitting, we tested if any of the three models were not a good fit for any of the areas of the anatomical or multi-modal levels. To test the goodness of fit for a given model and area, a deviance statistical test was performed. It assesses if the fit is significantly a better fit than a constant model. Multiple comparisons were FDR corrected 36. Areas that did not survive the multiple comparison correction were masked out and excluded from subsequent analyses.

### Subsampling analysis

We used a subsampling method to estimate developmental trajectories using a cross-sectional sample. This method is adapted from Vakorin et al., (2017) and Kozhemiako et al., (2018), and was used to statistically assess group differences in the trajectory shape captured by the highest order coefficient of a model between ASD and TD populations. In the present study, this subsampling method was applied to statistically assess if trajectory shapes in CT constant across age (slope, curvature or aberrancy) are atypical in the ASD group.

This method consists in generating small samples (subgroups) of subjects randomly pulled from the main group. This procedure is repeated 100,000 times, and a set number of sub-groups with the flattest age distribution, measured as sample entropy, are selected. The number of subgroups was set to 80. For each experimental group, 80 subgroups each composed of 70 subjects were selected. The number of subjects repeated across subgroups or the entropy (age distribution) between ASD and TD groups were not significantly different.

Once subgroups were created, the three models were fitted in all areas of each spatial level (HP, FSAP, MMP) where the model was proven to be a good fit as tested by a deviance test. From the three models, the coefficients termed slope, curvature and aberrancy were obtained, thus for each spatial level, a matrix of (80 subgroups × n areas × 3 coefficients) separately for ASD and CTR groups and were saved for the statistical multivariate analysis.

### Multivariate statistical analysis

To assess group differences in developmental trajectories, as well as to test for potential associations with ASD symptomatology, we applied Partial Least Squares (PLS) analysis. PLS is a multivariate statistical approach that uses singular value decomposition to extract Latent Variables (LVs), composed of a singular vector, singular value and LV design, explaining the variance in the data ^38^. In this study, two types of PLV were used: mean-centered and behavioral. The former is suitable for testing for an overall contrast between groups or conditions. In this study, a contrast expresses ASD > TD differences in trajectory shapes in CT. In contrast, the behavioral PLS explores the correlation between features and responses or behaviors, in our case, trajectory shapes and ADOS scores. Both approaches, perform one single permutation test by resampling without replacement subjects’ assignment across groups, and a unique p-value for all the features is obtained by counting the number of times a permuted singular value was higher than the original. Then, bootstrapping is performed by resampling with replacement subjects while keeping the group assignment fixed. The original singular vector is divided by the bootstrap standard error to obtain a bootstrap ratio associated with each feature indicating their reliability, akin to z-scores. In this paper, bootstrap ratios will be called z-scores. Each feature, in our case a coefficient expressing a trajectory shape in a given area from a parcellation, is associated with a z-score which indicates the stability of the area in reflecting the group differences. This procedure is commonly applied in statistical analysis across many fields, but especially in neuroimaging (McIntosh & Lobaugh, 2004; Krishnan, Williams, McIntosh, & Abdi, 2011).

### Analysis of group differences in trajectory shapes in CT

Three independent mean-centered PLS analyses were conducted at the hemispheric, anatomical and multimodal levels to detect differences in trajectory shapes between ASD and TD groups. For each PLS, the input was a matrix of size (80 subgroups × n areas × 3 coefficients) separately for ASD and CTR groups. Permutations and bootstrapping were performed 10,000 times to obtain statistically reliable results. Assuming the contrast ASD>TD, positive z-scores are associated with brain regions wherein the trajectory shape in CT is increased in the ASD group. Correspondingly, negative z-scores indicate brain regions with decreased CT trajectory shape in ASD. The z-scores for each model were separated, and each had a skewed distribution, indicating that there is a predominant increased or decreased effect representing the group differences. To visualize the results, each model coefficient was plotted separately and thresholded with the highest value opposite of the skewness.

### Center-specific analysis of group differences in trajectory shapes in CT

The same subsampling analysis of group differences was repeated for a single center, using the NYU repository. The number of subjects was reduced to 40 and the number of subgroups to 50. We aimed to obtain similar results while fully avoiding effects associated with inter-center variability.

### Associations between ADOS scores and trajectory shapes in CT

To explore the relationship between ASD symptomatology and trajectory shapes of CT maturation, behavioral PLS correlated the trajectory shape from a given model fitted in a subgroup with the subgroup mean ADOS-Generic scores (communication, social and stereotyped behavior subscales). Given that the trajectory shape of each model might correlate differently, for each of the three parameters (slope, curvature and aberrancy) at anatomical and fine-coarse spatial levels we performed three PLS (6 in total), and the p-values obtained for each model were Bonferroni corrected.

The behavioral PLS analysis renders an LV design vector indicating the contribution of the ADOS sub-score (communication, social and stereotyped behavior), a correlation for each ADOS sub-score, a single p-value for the overall-correlation, and z-scores expressing the stability of the feature in expressing the correlation. To make contributions of individual subjects more pronounced, the number of subjects per subgroup was decreased to 20, and to increase the combination of subjects, the number of subgroups was increased to 400.

### ASD classification of CT developmental trajectories

To assess which model parameter can better predict a developmental change from ASD or TD populations, a linear kernel support vector machine (SVM, using Matlab 2018b *fitcsvm* function) was trained to classify trajectory shapes from ASD and TD subgroups using the FSAP or MMP areas.

First, experimental groups from the full dataset were split into two equal training and testing sets. To have similar age, for each experimental group subjects were separated within two years age bins, and the subjects in each bin were randomly split in two. Once the training and testing sets were created, for each set, the subsampling analysis was applied (the CT values from the FSAP and MMP areas using 80 subgroups were fitted using the three models and the three coefficients were obtained). Then, an SVM was trained using the training set to classify one model coefficient from the FSAP or MMP, and the testing set was used to measure the accuracy in classifying ASD and TD using the slope, curvature or aberrancy. The accuracy was the average of specificity (how many ASD subgroups were correctly classified) and sensitivity (how many TD were correctly classified). This procedure was repeated 500 times (500 cross-validations) to obtain a distribution of accuracy, specificity and sensitivity.

### Data and code availability statement

Data is available from the ABIDE repository. The codes for the analyses are publicly available at https://github.com/AdoNunes/CT_trajectory_features_ASD_2019

## Discussion

In this study, we investigated atypical developmental trajectories of cortical thickness (CT) in the ASD during childhood and late adolescence. Previously, age-related changes in CT were studied by fitting a linear, quadratic and cubic models. Instead, in this study, we focused on the highest order derivative coefficient (the slope, curvature and aberrancy) to obtain an age-constant parameter that has an interpretable geometrical shape. These “trajectory features” describe a shape of the developmental trajectory of CT that describe the entire age range of the ASD and TD groups. We explored these features at several levels of spatial coarseness using a hemispheric, anatomical and multi-modal atlas. Overall, our results indicate that in ASD trajectory features are atypical in frontal, parietal and midline areas, and are a robust feature to predict ASD development, especially with the curvature parameter from the quadratic model.

Effects with strong directionality differences in CT trajectory features between ASD and TD groups were found using the quadratic and linear models, which had skewed z-score distributions indicating lower curvature and higher slope values in ASD. In this context, a more positive slope reflects a linearly decreased rate of cortical thinning, while a more negative slope expresses decreased rate of cortical thinning in early childhood and accelerated thinning after mid-adolescence. The slope and curvature coefficients although having opposite signs, represented a similar effect and the spatial representation of the strongest z-score values greatly overlapped. The trajectory shape captured by the quadratic model significantly correlated with the subgroup-averaged ADOS scores, indicating that the higher the symptom severity of the subsample the more negative the curvature coefficient was. Areas with the most stable correlation of curvature coefficients with ADOS scores tended to also be the most stable in reflecting group differences in curvature values between ASD and TD. This supports the association between higher ASD symptom severity and a more negative CT developmental curvature.

Our results are consistent with other studies which revealed increased CT in ASD individuals during childhood and early adolescence ^3^, indicating smaller decrease of CT in this period. A study with a large sample size of N=3000, found that alterations in the CT are more pronounced in the adolescence period, exhibiting a peak in CT especially in frontal areas and lessen in frontal areas ^42^. A recent CT study using the ABIDE I also found increased CT in the ASD from early childhood to adolescence and equal or reduced CT in early adulthood in frontal and parietal areas, indicating decreased CT thinning during childhood and adolescence and accelerated thinning in late adolescence ^43^. Studies using an age-range similar to ours concluded that the cortex thins less in ASD compared to typically developing controls ^3,44^. In addition, the developmental trajectories of the TD group of our study are very similar to the ones described in the longitudinal study of Zielinski et al. 2014, however, our ASD trajectories differ with this study in the period of accelerated thinning where it starts later than in Zielinski et al.

The areas with the highest z-score from the slope and curvature coefficients include areas of the Default Mode Network (DMN) and the frontoparietal network (FPN). Several ASD studies have found alterations in the DMN and FPN involving atypical functional connectivity patterns ^45–47^, more variability in the spatial networks ^9,10^, and altered gyrification and structural network architecture in ASD ^12,48^. The DMN has been linked to theory of mind, social cognition and inner referential processes ^49–53^. One of the characteristics of the DMN is to be deactivated during an externally gated task ^54–56^, and DMN tends to be anticorrelated with the FPN which is a goal oriented network ^57,58^. It has been reported that less DMN deactivation is associated with more stressful performance during stimulus-driven tasks ^59,60^, and it has been found that in ASD task deactivation of the DMN is deficient probably underpinning sensory feedback feedforward alterations ^61,62^. Moreover, typical development of the DMN displays a pattern of more sparse and mainly local connections to a more long-range and cohesive network in typical development ^63,64^. Also, there is evidence that long-distance connections fail to develop during adolescence in ASD population ^65^. Our study converges with the aforementioned literature signaling alteration in areas of the DMN and some of the FPN, and indicates that typical cortical neural development is altered and might underpin alterations of structure-function relations in areas of the DMN and FPN.

The trajectory shape for CT in the TD group represents a neurotypical U-shaped thinning centered around the age of pubescence and early adulthood ^21,24^, whereas in the ASD group, there is a decreased thinning rate which is better fit by an inverse U-shaped curvature or a more positive slope using a linear model. This group difference is likely to reflect alterations in neural pruning associated with this age range. Neuronal pruning is the process of reducing the number of connections during the first two decades of life in order to select the most efficient and optimal connections, and evidence suggests that CT is likely to reflect dendritic arborization ^66^, Accordingly, maturational changes of CT most possibly reflect the process of dendritic pruning. It is currently understood that the number of dendritic synapses reach their peak during childhood and decrease during puberty ^67^. The hypothesis of altered neural, and in particular dendritic, pruning in ASD has become widespread as it is consistent with considerable evidence from multiple lines of research indicating brain overgrowth in ASD ^68,69^. It has been suggested that ASD is characterized by reduced synaptic pruning ^70,71^, and by genetic alterations impacting synaptic structure, function and regulation^72,73^. Our results, consistent with previous literature, give further support that atypical neural pruning in ASD is characterized by a decreased thinning rate of cortical thickness between childhood and adolescence and accelerated thinning during late adolescence.

Given that the examined trajectory shapes of CT across age are constant, speculatively these rates of CT change could be estimated from one subject having several longitudinal MRI acquisitions. Using a cross-sectional data, we obtained a good accuracy in classifying CT developmental features, especially using the curvature of the quadratic model and using the areas from the MMP atlas. This suggests that similar accuracy could be obtained when trying to predict if individuals have ASD based on their longitudinal CT measures. To estimate the curvature in the quadratic model is necessary at least three time points. Given that our results indicate that the quadratic model is the most accurate and sensitive parameter in classifying ASD, we suggest that three longitudinal time points would be necessary to predict if an individual has ASD.

## Abbreviations

(ASD): Autism Spectrum Disorder
(TD): Typically Developing
(CT): Cortical Thickness
(ADOS): Autism Diagnostic Observation Schedule
(PLS): Partial Least Squares.

## ACKNOWLEDGEMENTS

This study was supported by the Canadian Institutes of Health Research: CIHR to S. M. D. (MOP-136935)

## FINANCIAL DISCLOSURES/ CONFLICT OF INTEREST

These authors report no biomedical financial interests or potential conflicts.

**Supplementary Figure 1.**
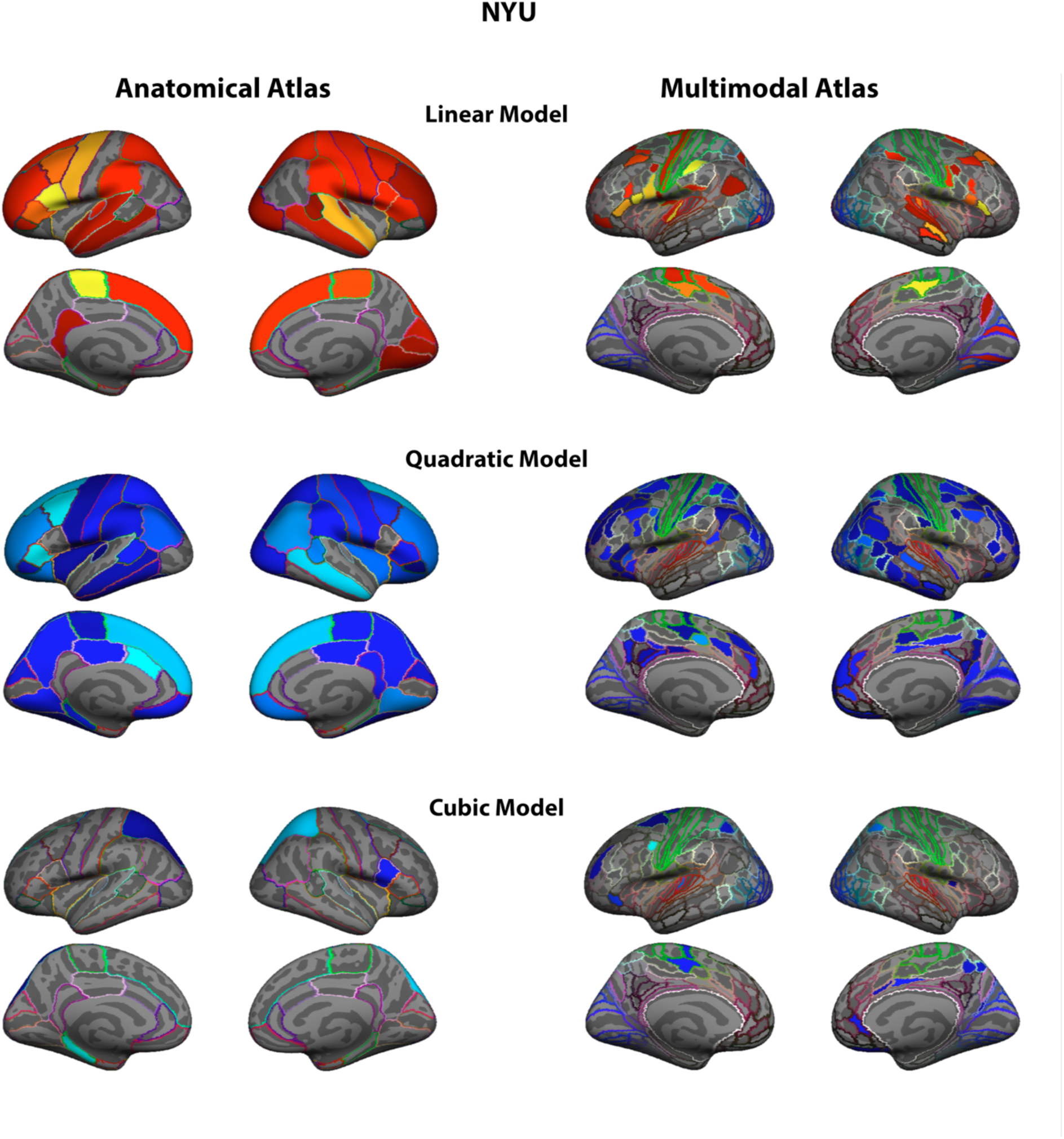
Group differences in rate of changes in CT maturation using the NYU center. For the group contrast (ASD>TD) the effect found in the linear coefficient is positive, whereas in the quadratic and cubic is negative. The spatial distribution of the highest z-scores in the linear model are very similar to the quadratic, and both represent a decreased cortical thinning during childhood to adolescence maturation but the quadratic also captures accelerated thinning in the later age period. On the left are the results from the anatomically defined FreeSurfer atlas and on the right the Multi-Modal Parceled atlas. Outlined colors in each cortical area represent the original atlas annotation color of the area.

